# Nesso-1: Accelerating Open-Source Binding Affinity Predictions

**DOI:** 10.64898/2026.08.01.742196

**Authors:** Nikhil Shenoy, David Errington, Emmanuel Bengio, Kacper Kapuśniak, Kerstin Klaeser, Yui Tik Pang, Vladimir Radenkovic, Prudencio Tossou, Therence Bois, Andrew Wedlake, Francesco Di Giovanni

## Abstract

In this technical report, we introduce Nesso-1, a *coarse-grained* cofolding framework for binding-affinity prediction. Nesso-1 requires *∼* 1 second per prediction on a single GPU. This offers more than one order of magnitude speed-up over the leading open-source baseline, Boltz-2, which significantly expands the regions of chemical space that can be explored during high-throughput virtual screening. Importantly, Nesso-1 matches or surpasses the accuracy of Boltz-2 over the same benchmarks adopted in their study—which we show reflect in-distribution scenarios—as well as over more challenging out-of-distribution data encompassing the OpenBind affinity benchmark and 25 internal biochemical assays. Notably, Nesso-1 maintains robust predictive accuracy even on assays with extremely low similarity to the training data. Moreover, we highlight examples where Nesso-1 demonstrates meaningful selectivity, separating the binding affinities of identical compounds between on-targets and related off-targets. Nonetheless, zero-shot generalization to real-world medicinal chemistry remains an inherently challenging task; consequently, we acknowledge specific assays where the model’s performance is limited. We **open-source** Nesso-1: code and weights are available at https://github.com/recursionpharma/nesso

## 1 Introduction

A key challenge in target-based drug design is predicting whether a compound binds to a protein and its associated *binding affinity*, which represents how energetically favourable such an interaction is [1]. For a given target, finding hits requires a scoring function that is reliable *at scale* across chemical space. However, in later stage design, *precision* plays an increasingly important role and understanding specific interactions in the binding site is necessary to identify the most potent and promising candidates. Since binding affinity derives from enthalpic and entropic contributions, estimating this value ideally involves knowing or predicting the 3D *structure(s)* that a protein-ligand complex can assume. However, explicit modeling of every atom in the 3D structure introduces a vast number of degrees of freedom and has resulted in unforgiving trade-offs between speed and accuracy. Docking methods have been a primary workhorse in structure-based drug design thanks to their efficiency [2; 3; 4]. Yet their accuracy greatly depends on access to a suitable *holo* structure, as they may not capture induced-fit conformational changes [5; 6]. Alternatively, simulation-based methods such as Free Energy Perturbation (FEP) [7; 8; 9; 10] have shown high correlation with experiments [11; 12] as they rely on molecular dynamics to capture thermodynamic changes. However, such approaches incur a high computational cost [13] and are sensitive to structure preparation and simulation protocol [14].

**Cofolding** methods, whereby deep-learning models directly predict the holo 3D structure of a protein-ligand pair from sequence and SMILES, have shown promise in disrupting the physics-based Pareto front [15; 16; 17; 18; 19; 20; 21; 22; 23]. A long-term aspiration is to benefit from these methods not only to predict structures but also to score their affinity accurately across diverse targets and chemotypes. The main cofolding paradigm relies on training an *expensive* trunk that updates single and pair representations, which are then refined by a heavy-atom generative model. Boltz-2 expanded upon this framework by leveraging both the trunk outputs and the generated poses to train a separate affinity module [24]. As the *leading open-source* cofolding affinity model, Boltz-2 demonstrates that predictive accuracy significantly surpassing traditional docking scores can be achieved without the need for computationally expensive simulations. However, subsequent evaluations have underlined limited out-of-distribution performance [25; 26]. Importantly, Boltz-2 affinity predictions can be uncorrelated with the quality of the generated poses [27; 28], suggesting that atomistic resolution may be unnecessary or even harmful for the affinity module. In fact, such resolution means that a prediction requires *∼* 20 seconds, a cost which remains prohibitive in large Hit-ID campaigns. In light of this, we raise the following:

**Key Problem.** Full-atom resolution in cofolding models severely limits applicability in early drug-discovery stages and may be unnecessary when training a separate binding affinity module.

Faster alternatives to Boltz-2 exist, yet these are achieved by effectively removing cofolding altogether. Methods like Hermes [29] directly predict affinity without explicitly modeling structures; while this approach offers orders of magnitude speed-up, its transferability across protein classes is unclear. Boltzina mitigates the cost of sampling from a generative model, but this derives from reverting to rigid docking [30]. Ligand-only generative models often excel at pose prediction yet they require knowledge of the binding site and may ignore the receptor’s flexibility [31; 32; 33]. While these approaches are generally viable, it remains unclear to what extent removing structural information or keeping the target rigid is sufficient for downstream predictions.

TerraBind is a recent model which tackles the **Key Problem** by operating on the coarse-grained token-level representation only and removing the heavy-atom generative model [34]. The affinity module is instead optimized exactly as in Boltz-2. Thanks to these changes, TerraBind reports to be one order of magnitude faster than Boltz-2 and achieve comparable or superior accuracy in affinity prediction. Although this work represents a significant advance, important limitations persist: (i) TerraBind is *closed-source* and depends on *proprietary* ligand embeddings; (ii) Evaluation on public benchmarks is limited (CASP16 only) and it remains difficult to assess similarity between their assay chemistry and the data that TerraBind and the proprietary embeddings were trained on.

To address both the **Key Problem** and TerraBind’s limitations, in this report we present Nesso-1, a coarse-grained cofolding model for predicting binding affinity. Nesso-1 adopts the training data and\ optimization framework of Boltz-2 [24] but removes much of its architectural complexity, in particular the atomistic Diffusion Module, in line with the valuable changes proposed in TerraBind [34]. In our report, we primarily compare Nesso-1 against Boltz-2, as this is the leading *open-source* cofolding-based affinity model and utilizes the same affinity training data. Our *main contributions* are:

- Nesso-1 **is open-source and does not rely on proprietary ligand embeddings**. We release weights trained strictly on public data, providing the broader community with a fast, accessible cofolding affinity model that can be deployed at scale for high-throughput screening.
- Nesso-1 **is the fastest open-source cofolding affinity model.** Requiring *∼* 1 second per prediction, Nesso-1 is *>* 10*×* faster than Boltz-2 (see Figure 1, left). Internally, this speed-up allows us to screen up to 20*×* the number of compounds compared to Boltz-2 under the same computational budget. Consequently, Nesso-1 significantly expands the breadth of high-throughput virtual screening at the accuracy of cofolding affinity models and facilitates rapid fine-tuning on continuous data streams.
- Nesso-1 **matches or surpasses the accuracy of leading open-source affinity models.** Nesso-1 outperforms Boltz-2 on their original benchmarks. As we demonstrate that these datasets share high chemical similarity with the training data, we further establish Nesso-1’s robustness in challenging, *out-of-distribution* scenarios. Nesso-1 achieves the highest accuracy on the recent OpenBind benchmark [6] and outperforms Boltz-2 on 25 internal, biochemical assays (see Figure 1, right).
- Nesso-1 **remains competitive out-of-distribution**. We show that Nesso-1’s predictive accuracy is robust across varying degrees of chemical similarity to the training data. Moreover, we provide examples where Nesso-1 has meaningful *selectivity* for detecting off-target liabilities.

**Figure 1:**
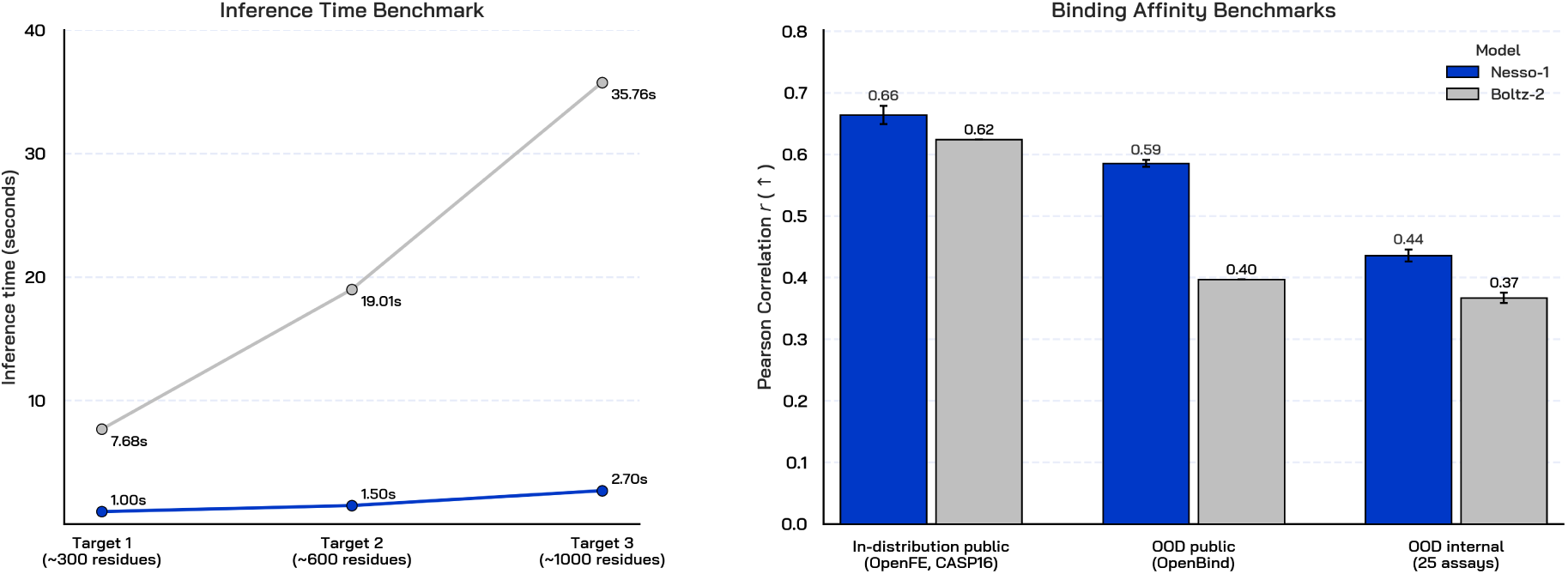
Nesso-1 accelerates open-source cofolding affinity models (Boltz-2) while matching or surpassing their accuracy. On the left we compute inference runtime over three internal targets: estimates are obtained on a single H100, averaging over 10 compounds, and do not include fixed costs (ESM-2 loading for Nesso-1 and MSA for Boltz-2). Note that both models leverage cuEquivariance kernels. On average, Nesso-1 is *>* 10*×* faster than Boltz-2. On the right, we report cross-assay Pearson averages weighted by the number of compounds. Error bars are computed across recycling steps for Nesso-1 and for Boltz-2. For Boltz-2, error bars are only reported for evaluations conducted in this study. We distinguish between in-distribution and out-of-distribution benchmarks based on the chemical similarity with the training data. Nesso-1 outperforms Boltz-2 across each benchmark. In general, zero-shot generalization to real-world medicinal chemistry is a challenging task and the overall performance of both models reflect that.

**How to interpret our results** Figure 1 illustrates the contrast between standard public benchmarks and real-world biochemical assays. Specifically, our report highlights that test sets like OpenFE and CASP16 are largely in-distribution for models trained on BindingDB or ChEMBL. Consequently, we caution against drawing broad comparisons between Nesso-1 and Free Energy Perturbation (FEP) methods based solely on these benchmark performances. Indeed, zero-shot generalization to real-world medicinal chemistry for models trained exclusively on (biased) public data remains an exceptionally challenging task, as similarly noted by [33]. It is therefore expected that the performance of Nesso-1 and Boltz-2 is limited on certain internal assays. Indeed, even on the recent OpenBind benchmark, Nesso-1 offers only a modest improvement over a simple molecular weight ranking. Nevertheless, Nesso-1 surpasses existing AI models on OpenBind by a notable margin and remains competitive on most internal assays featuring heavily out-of-distribution chemistry. Crucially, it manages to outperform the leading open-source baseline while offering more than an order of magnitude speed-up.

*Despite limitations in zero-shot generalization on certain assays,* Nesso-1 *currently establishes a faster cofolding affinity baseline that matches or surpasses the accuracy of leading open-source models. We release it as a foundational tool for the community to build upon and iteratively improve*.

## 2 Results

### 2.1 Overview of model and data

In this technical report we mainly focus on empirical evaluations. Before discussing the results, we provide an outline of Nesso-1’s architecture and training (see Figure 2). Nesso-1 adopts the following general principles:

i. Removing the heavy-atom diffusion module as proposed by TerraBind [34] and streamlining the trunk to only update pair-token representations [35; 34; 36];
ii. Curating public affinity data as reported in Boltz-2 [24] and employing similar losses that focus on intra-assay differences to compensate for noise and biases in different experimental settings.

**Figure 2:**
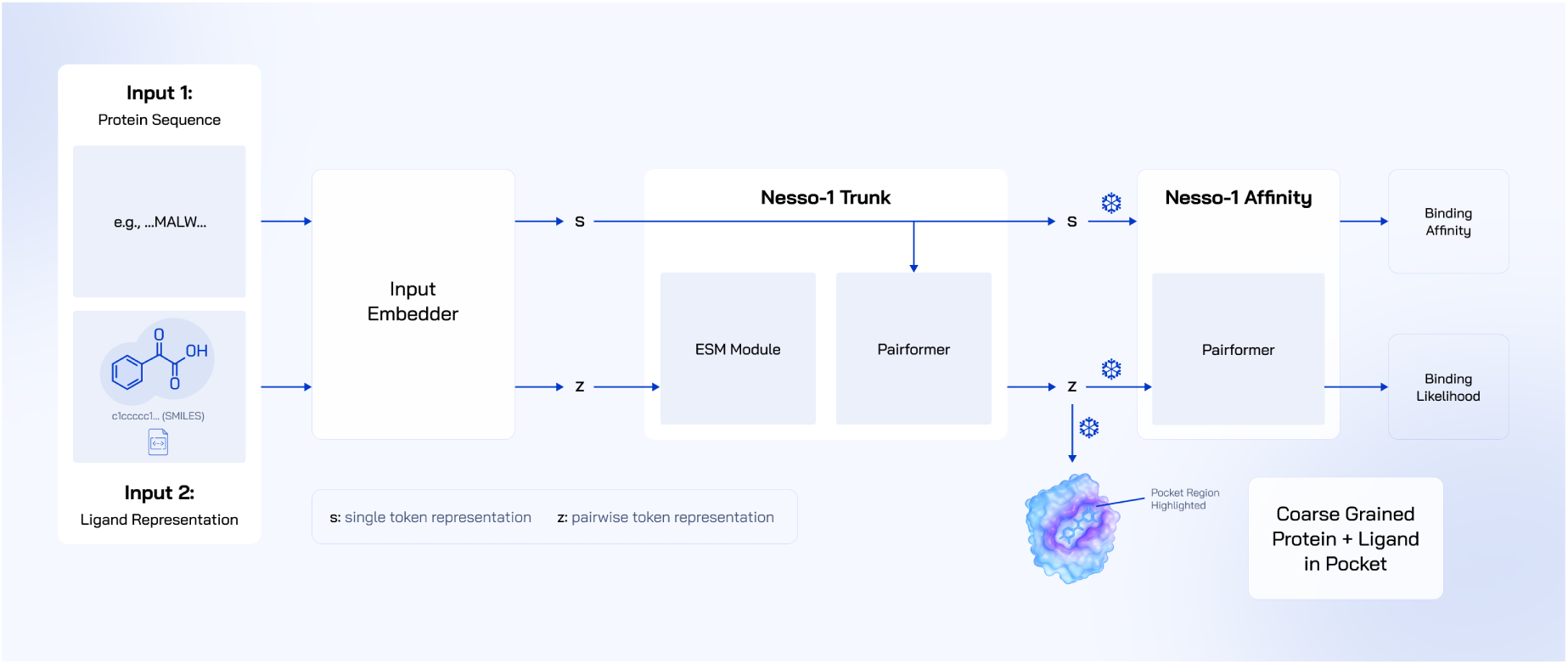
Diagram of Nesso-1’s architecture.

While TerraBind adopts a similar paradigm [34], it relies on proprietary ligand embeddings and remains closed-source. In contrast, with Nesso-1 we release weights that are independent of proprietary embeddings and trained solely on public data. Our results show that Nesso-1 not only offers significant speed-ups compared to leading open-source baselines such as Boltz-2 but often exceeds their accuracy. These findings indicate that existing structure-based models may fail to map specific atomistic interactions to binding-affinity, and hence may be limited by such high-frequency structural information.

#### Structure module

Nesso-1 follows a general cofolding blueprint, where a sequence and a SMILES are first embedded via an *atom encoder* and then mapped to a pairwise representation *z* by a Pairformer *trunk* [16; 21]. The latter relies on triangle attention and multiplication operations instantiated at the token-level, meaning that *z* directly encodes geometric information regarding ligand heavy atoms and protein residue centers (*C_β_*, or *C_α_* in the case of Glycine). In line with later simplifications proposed in [35; 34], the trunk in Nesso-1 only updates pairwise token representations, resulting in a significant reduction in parameters. Moreover, we also eliminate MSA information following [34; 36]. To compensate for a more efficient trunk with less capacity, we rely on pre-trained protein embeddings from ESM-2 [37].

The representation *z* is optimized via categorical cross-entropy over distance bins. We use the center of these bins and the predicted logits from the trunk to infer the *expected distance* between tokens. From the probability distribution associated with these pairwise distances, we can compute a normalized entropy restricted to protein-ligand pairs only, which we denote by *H_P L_* as in [34]. During inference, expected distances are used for cropping and to reduce context for the affinity module, while the protein-ligand entropy *H_P L_* acts as an intrinsic structure uncertainty module. In particular, we adopt TerraBind’s cropping strategy, whereby the full protein context is used only during the first pass of the trunk, while the remaining recycling steps retain protein tokens with predicted distances ≤ 22Å from the ligand.

#### Training data and stages

The trunk and atom encoder are trained on *public data only*. In particular, we leverage experimental crystal structure from PDB released before 2021-09-30 [38], high-quality protein distillation data from AFDB [15; 39] and a subset of distilled protein-ligand complexes from SAIR [40] curated and released by [33]. Training follows a multi-stage strategy aligned with [34], where we progressively focus on the binding site through cropping.

#### Affinity Module

As proposed in Boltz-2 [24], we instantiate a separate, smaller Pairformer, termed the *affinity module*, which is optimized to predict both a binding likelihood value (for Hit-ID classification) and a binding affinity value (for Hit-to-Lead regression). We treat binding affinity and potency labels (*K_i_, K_d_, IC*_50_*, EC*_50_) interchangeably to absorb the experimental biases and missing metadata inherent to public assay datasets. The affinity module conditions on (i) the pre-trained atom encoder, (ii) the frozen protein language model embeddings and (iii) the pre-trained pairwise representation *z*, alongside its inferred distance bin information. In particular, we follow TerraBind [34] and mask *z* so as to only include protein tokens within an expected cutoff of 15Åfrom any ligand heavy atom as predicted from the trunk.

#### Training data and optimization

The data curation follows *exactly* what is reported in Boltz-2 [24], which also allows for a fair comparison of models. Namely, we consider the same data as in [24, Table 1]. The main sources for Hit-to-Lead assays are BindingDB [41] and ChEMBL (v34) [42]; for classification data, we use PubChem (1.8.1) [43], CeMM Fragment Dataset [44] and the MIDAS Metabolite Data [45]. We employ the same decoy generation strategy, exclude the same uninformative assays, and apply identical data-leakage constraints by removing any training datapoint with *≥* 90% protein sequence similarity to the FEP+ benchmark [11]. A key distinction, however, is *how* structure-based filtering is implemented in practice. Adopting the heuristic from TerraBind [34], we discard any complex with protein-ligand entropy *H_P L_ >* 0.7 and pIC50 *≥* 6 (recall that pIC50 = *−* log *IC*_50_). The affinity module is jointly optimized using a focal loss for classification, forcing the model to learn to separate active compounds from inactive ones, and a Huber loss [46] for regression. The regression loss is explicitly split into an absolute loss and a relative difference loss, with the latter being up-weighted to focus the model’s capacity on capturing *intra-assay* differences and mitigate the effect of *inter-assay* biases [24].

The final weights are obtained by averaging the final training checkpoint and the best one from the validation loss.

### 2.2 Structural benchmarks

First, we evaluate the structural quality of the pairwise representation *z* learned by the trunk, which serves as the core input to the affinity module. To generate 3D coarse-grained poses, we adopt the optimization routine introduced by [34]. Here, the *C_β_* (*C_α_* for Glycine) and ligand-heavy atom positions are iteratively updated so that their spatial distances match the expected values derived from the distogram induced by *z*. We also benchmark Nesso-1_full_*_−_*_context_, where the full-protein context is leveraged at each recycling step (not just the first one). We adhere to the same evaluation criteria of TerraBind to test Nesso-1’s structural representations, which do not depend on proprietary ligand embeddings. All baseline performance numbers are taken from [34]—in particular, we recall that Boltz-1 has the same training data cutoff. For a fair comparison, we follow previous benchmarking efforts and also generate 10 pose samples per complex and select the top-scoring pose for evaluation after applying symmetry correction. Note that we adopt the same structural metrics: (i) Ligand RMSD < 2 Å success rate—after aligning over a region of 15Å centred at the ligand; (ii) Ligand RMSD < 2Å and LDDT-PLI *>* 0.8, which instead is enforced on a more localized region of 6Å. Our benchmarks include FoldBench [47], PoseBusters [48], and Runs N’ Poses [49]. Finally, minor discrepancies may exist between our evaluation sets and those of TerraBind due to unshared filtering criteria (e.g., specific small-molecule exclusions).

#### Analysis of results

The evaluation in Figure 3 demonstrates that Nesso-1_full_*_−_*_context_ matches or slightly surpasses the performance reported by closed-source models like TerraBind. Crucially, this is achieved without the need for proprietary ligand embeddings. While the performance of Nesso-1 on structural metrics is marginally worse than Nesso-1_full_*_−_*_context_, restricting the full-protein context solely to the first recycling step yields significant acceleration, particularly for large proteins. Consequently, in the following, we benchmark affinity predictions derived from the Nesso-1 trunk output. Indeed, internal evaluations reveal no noticeable gain in binding-affinity accuracy when utilizing the more computationally expensive representations from Nesso-1_full_*_−_*_context_. We acknowledge that heavy-atom resolution via generative modeling may still be necessary for precise cofolding, as our model cannot resolve exact side-chain placements within the binding pocket. Nevertheless, our results highlight that the representations learned by a streamlined trunk can be sufficiently expressive to capture the coarse-grained protein-ligand interfaces required for accurate binding-affinity predictions.

**Figure 3:**
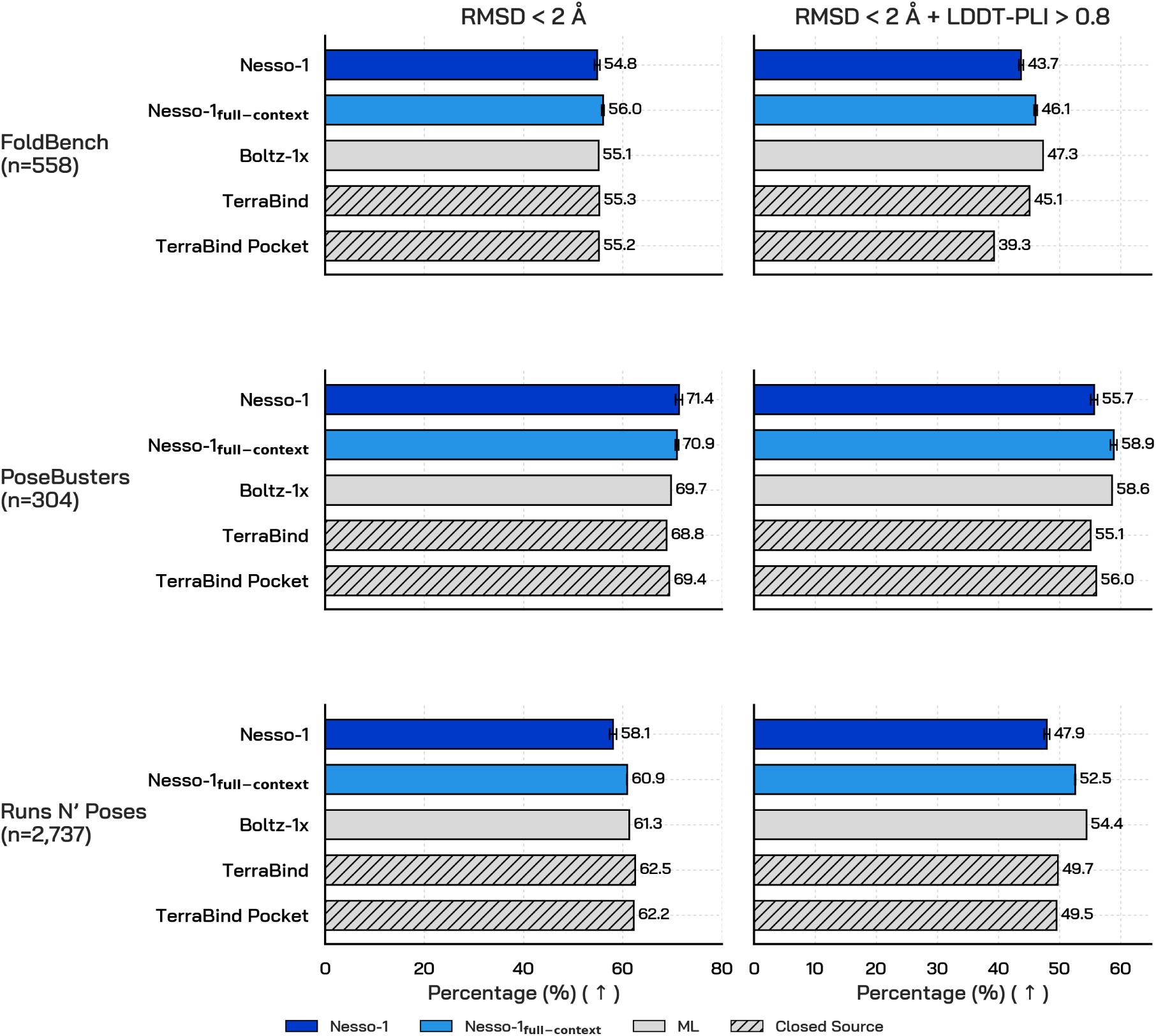
Evaluation of coarse-grained poses of Nesso-1 against baselines reported in [34]. Nesso-1 matches the performance of similar *closed-source* coarse-grained models such as TerraBind.

### 2.3 Binding-affinity benchmarks

The focus of our evaluations is on binding-affinity predictions. We include both public benchmarks and internal biochemical assays. We reiterate that Nesso-1 was trained on public affinity data following the curation strategy outlined in Boltz-2 [24], allowing for a fair comparison. Indeed, we primarily compare the speed-accuracy trade-off of Nesso-1 with that of Boltz-2, as the latter represents the leading open-source cofolding affinity model currently available.

#### 2.3.1 Inference speed

We benchmark the speed-up gains afforded by the coarse-grained cofolding approach in Nesso-1 by comparing with Boltz-2. In Figure 5 we consider three internal targets of varying sizes, ranging from *∼* 300 residues to *∼* 1000 residues. The wall-clock inference time of each model is an average over 10 compounds per target. In particular, the estimates account for the cost of running each model from input to binding-affinity predictions, but discarding fixed per-target calculations like loading protein embeddings for Nesso-1 and running MSA for Boltz-2, respectively. Our benchmarking demonstrates that Nesso-1 is, on average, one order of magnitude faster than Boltz-2, with speed-up surpassing 20*×* on larger proteins. In particular, even though both models benefit from NVIDIA cuEquivariance kernels [50], the acceleration is even more significant in the case of Nesso-1. By drastically reducing computational overhead, Nesso-1*unlocks the ability to perform high-throughput screening efficiently and renders large-scale fine-tuning accessible for downstream applications*.

After demonstrating that Nesso-1 is the fastest open-source cofolding affinity model, we next focus on evaluating its accuracy for affinity predictions. We first test Nesso-1’s ability to predict binding likelihood and binding-affinity values on the public benchmarks originally adopted in the Boltz-2 paper [24]. Additionally, we run Nesso-1 zero-shot on the recent OpenBind benchmark, which arguably represents a more challenging and realistic example of a Hit-to-Lead stage in a drug-discovery campaign.

#### 2.3.2 MF-PCBA benchmark

We assess the performance of the classification head in separating active compounds from inactive ones. This is a task that is particularly relevant during Hit-ID stages in a drug-discovery campaign. To this aim, we evaluate Nesso-1 on the MF-PCBA benchmark [51], a collection of datasets including up to 16 million unique protein-ligand interactions, which is aimed at emulating large high-throughput screening in a campaign. We note that this benchmark was originally adopted in the Boltz-2 work [24], from which we directly source the baseline metrics for models such as GAT, BACPI [52], and Chemgauss4 [53]. As shown in Figure 4, we report the average precision and enrichment factors at top-ranked percentiles, consistent with prior work. Our empirical results demonstrate that Nesso-1 matches or exceeds the accuracy of its all-atom open-source counterpart (Boltz-2), while outperforming the other docking and machine learning baselines by a significant margin.

**Figure 4:**
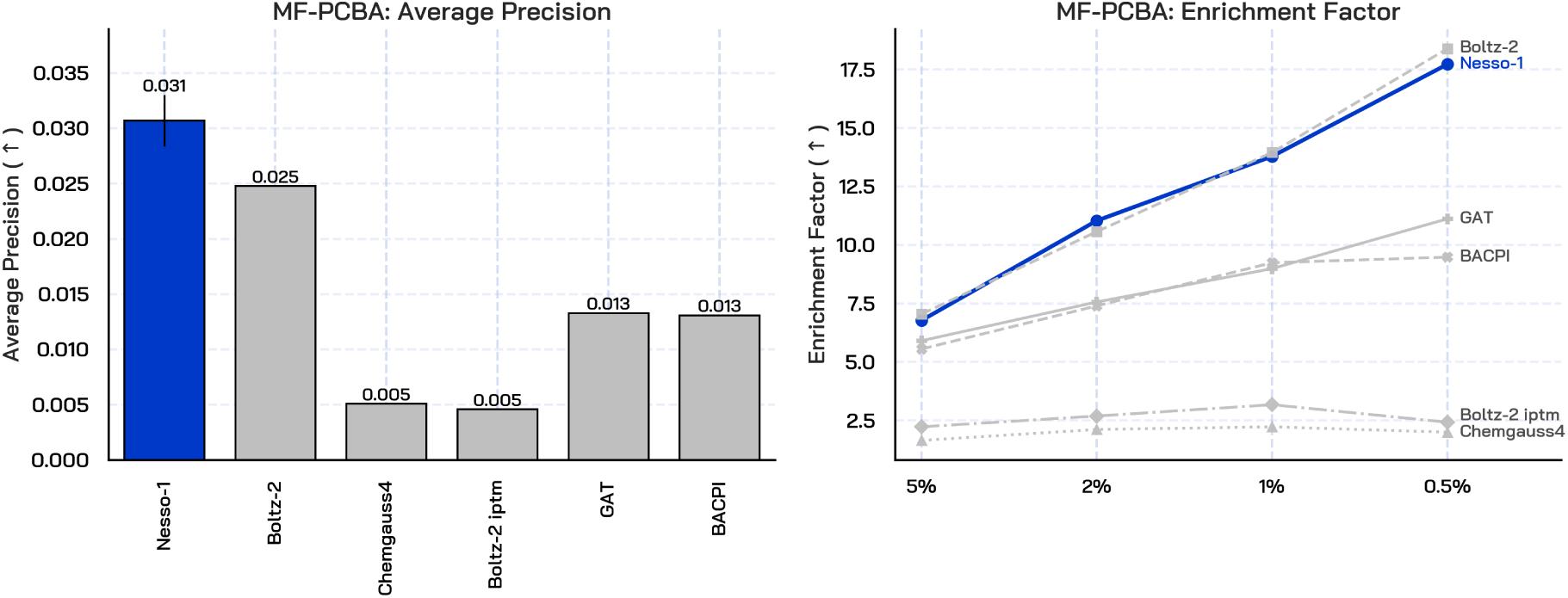
Average Precision and Enrichment Factor at different percentiles on the MF-PCBA dataset, as measured from the predictions of the classification head in Nesso-1’s affinity module. Results confirm that Nesso-1 achieves the highest average precision and matches Boltz-2’s enrichment factor.

**Figure 5:**
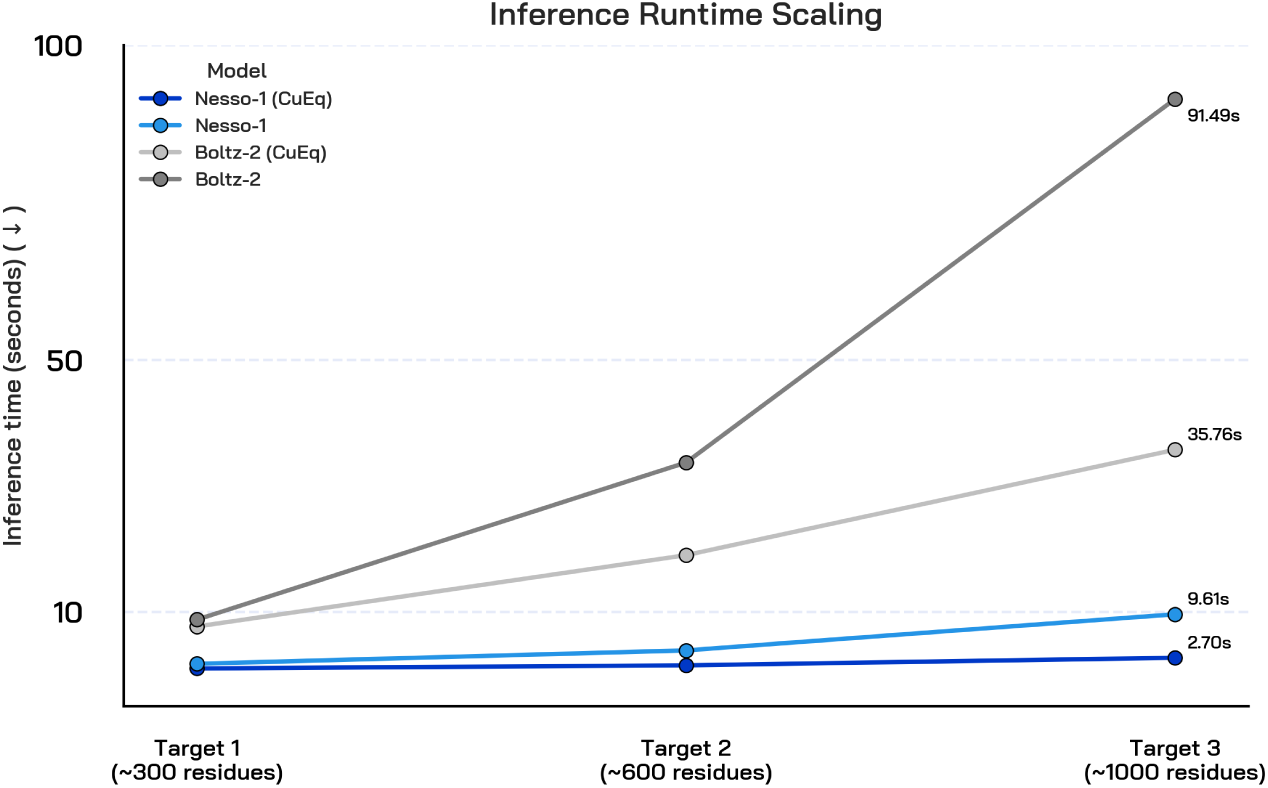
Inference runtime of Nesso-1 and Boltz-2, both tested with and without NVIDIA cuEquiv- ariance kernels [50]. We compute inference runtime over three internal targets: estimates are obtained on a single H100, averaging over 10 compounds, and do not include fixed costs (ESM-2 loading for Nesso-1 and MSA for Boltz-2). On average, Nesso-1 is *>* 10*×* faster than Boltz-2. While both models benefit from cuEquivariance kernels, the acceleration is even greater in the case of Nesso-1.

#### 2.3.3 FEP+ and CASP16 benchmarks

We evaluate the performance of Nesso-1 on public benchmarks adopted by Boltz-2 that simulate the Hit-to-Lead and Lead-Optimization stages of a campaign. In these scenarios, the primary objective is to identify which compounds in a given series exhibit the highest binding affinity. First, we consider two subsets from the FEP+ benchmark [11], namely the OpenFE dataset [54] and a smaller subset of 4 targets where we have access to Absolute Binding Free Energy (ABFE) calculations [10; 55]—a free-energy-perturbation method relying on running multiple molecular-dynamics simulations per protein-ligand pair. The OpenFE dataset includes 876 protein-ligand pairs. Conversely, the FEP+ 4 subset has 87 neutral compounds across 4 *kinases*: CDK2, TYK2, JNK1, P38. We also evaluate on the CASP16 affinity benchmark [56], which spans 140 complexes across two targets.

In addition to Boltz-2, we assess the performance of Nesso-1 against computationally intensive simulation-based methods such as FEP+, ABFE, and OpenFE. We emphasize that the results for any such baseline are sourced directly from [24], which also provides the performance of cheaper but less accurate physics-based methods such as Fragment Molecular Orbital and MM/PBSA. Furthermore, on CASP16 we also report the performance of TerraBind [34], the closed-source counterpart of Nesso-1, as this is the only public affinity benchmark evaluated in their study. Our final baseline is IsoDDE [57], another closed-source model where comparison is significantly more difficult due to the lack of architectural and training details. We report the average Pearson correlation across assays weighted by the number of compounds—as originally done in [24]. This choice reflects the fact that, in practice, we are often more interested in comparing the affinity (or potency) of compounds in the *same* assay rather than predicting an exact value. Moreover, weighting by the number of compounds, we avoid outliers produced by small assays. Finally, while IsoDDE’s performance is taken from its technical report, the lack of methodological details regarding their Pearson correlation calculation introduces potential comparative discrepancies.

##### Analysis of results

Figure 6 shows that Nesso-1 outperforms the state-of-the-art, open-source cofolding affinity model, Boltz-2. The results further confirm that removing heavy-atom resolution does not hurt binding-affinity prediction accuracy, when compared to existing baselines. In fact, Nesso-1 even surpasses the performance of simulation-based approaches such as OpenFE and ABFE. Note that these take 6-12 GPU hours [24] for a single prediction, making Nesso-1 more than 10^4^*×* faster. While these results offer a controlled way of comparing cofolding models against physics baselines, limitations of these benchmarks—discussed below—prevent us from drawing universal conclusions. Nevertheless, it is equally not straightforward to extrapolate the performance of Free Energy Perturbation protocols beyond these benchmarks, as their accuracy greatly depends on structure preparation, simulation parameters, and protein class [58; 14]. Finally, we assess the performance of Nesso-1 against *closed-source* models. On CASP16, Nesso-1 is comparable with the performance reported by TerraBind—note that this dataset only includes two targets and a total of 140 complexes. Conversely, IsoDDE’s reported performance is the highest on each benchmark. Crucially, in the following paragraph we detail several key limitations of these benchmarks, suggesting that absolute performance metrics and exact model rankings should be interpreted with caution. A concurrent work highlights this very same point [59].

**Figure 6:**
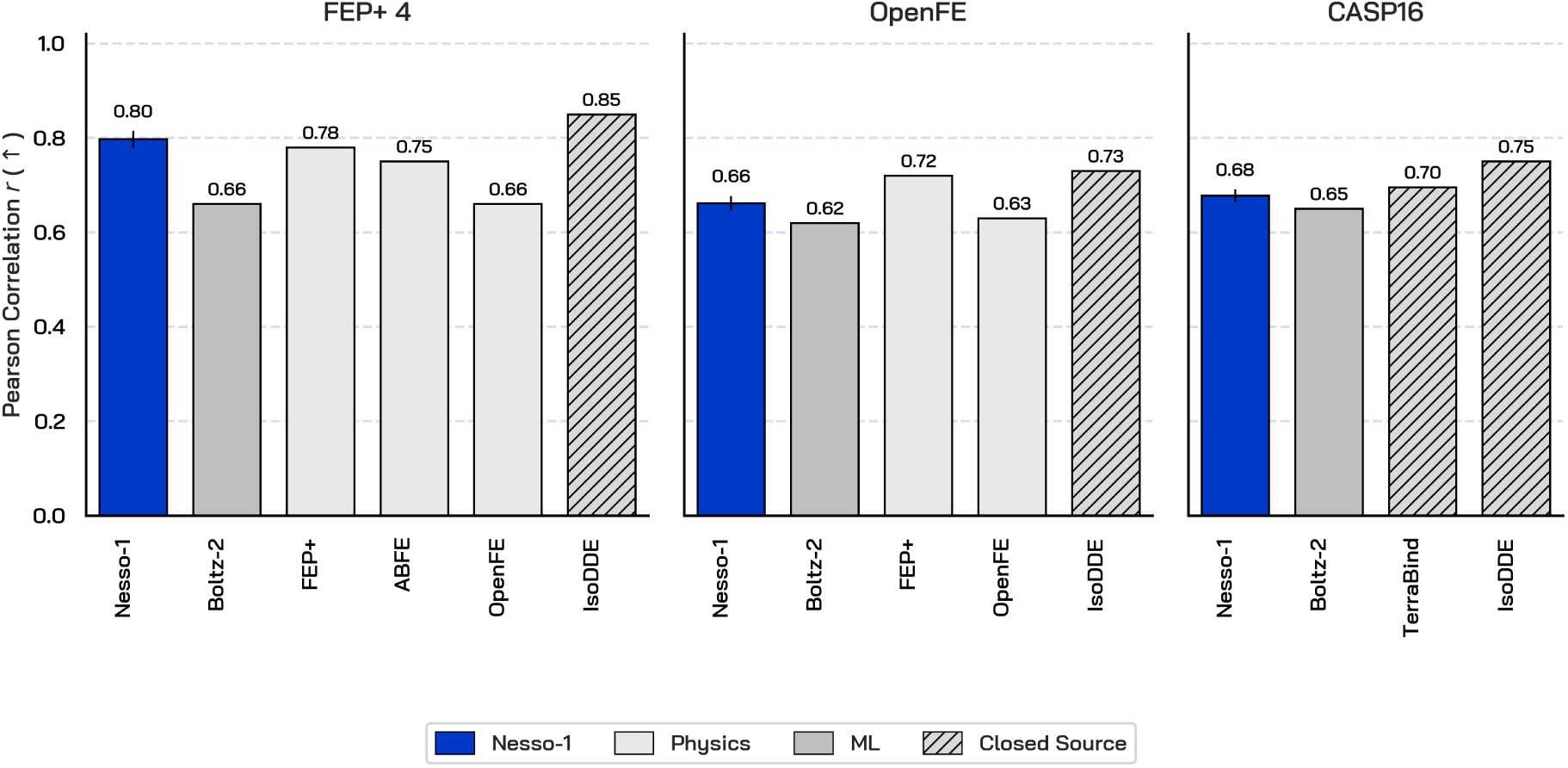
Average Pearson Correlation across assays weighted by number of compounds—note that IsoDDE does not share if the reported correlation is weighted. We compare Nesso-1 against open-source models (Boltz-2), closed-source models (IsoDDE, TerraBind)—*where available*—and physics-based methods relying on Molecular Dynamics (FEP+, ABFE, OpenFE). Results show that Nesso-1 is the most accurate open-source, cofolding affinity model. However, we emphasize that these benchmarks are, largely, in-distribution.

##### Limitations of these benchmarks

Nesso-1 and Boltz-2 train on the same data and use the same leakage constraints where proteins with sequence similarity higher than 90% to any target in the FEP+ benchmark are removed from the training split. At a high level, IsoDDE reportedly follows the same protocol, while no exact details have been disclosed by TerraBind. While adopting the same approach allows for a controlled comparison among baselines, particularly open-source models, the FEP+ and CASP16 splits *should not be regarded as indicative of performance on real-world medicinal chemistry*. Indeed, in Figure 7 we illustrate that these benchmarks have high chemical similarity to the public training data. Internal assays as well as the latest OpenBind benchmark are significantly more out-of-distribution and hence represent more challenging and realistic evaluation scenarios. Importantly, we have found that these in-distribution benchmarks typically reward overfitting. Indeed, we can produce checkpoints that achieve higher accuracy on each dataset by generally training for longer and removing any entropy-based filtering. However, doing so comes at the expense of generalization as it results in a model that significantly underperforms on harder assays. This finding aligns with concurrent work in [59], which provides a detailed overview of the limitations of evaluating models trained on BindingDB or ChEMBL when only sequence-similarity leakage constraints are enforced.

**Figure 7:**
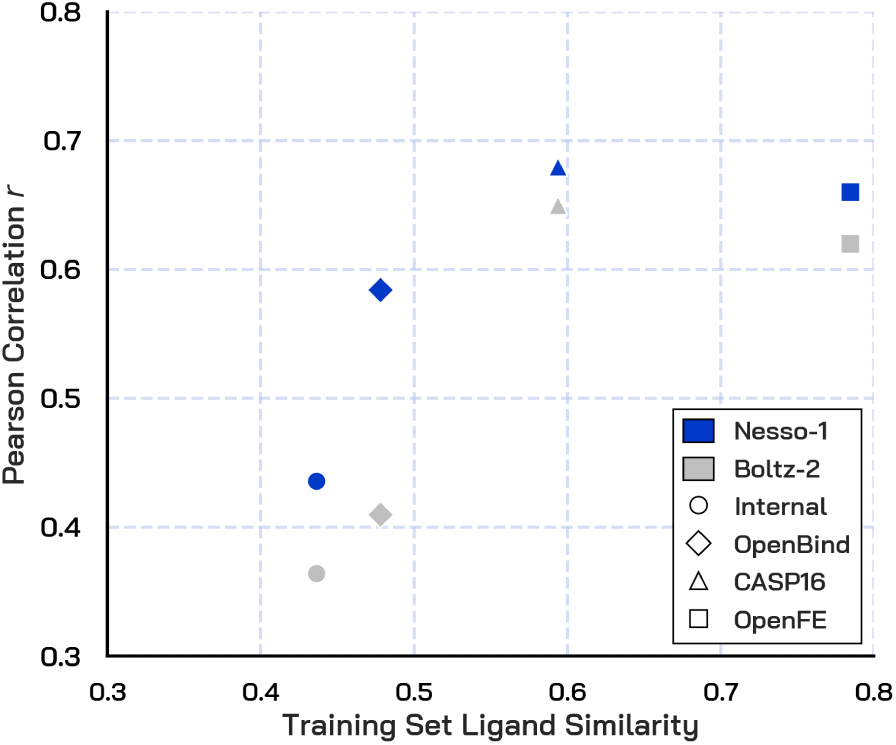
Pearson Correlation of Nesso-1 and Boltz-2 as a function of the chemical similarity to the affinity module’s training data. For a given target, similarity is defined as the mean maximum Tanimoto similarity (computed using Morgan fingerprints) between the test ligands and the training set. The final dataset-level similarity is obtained by taking the mean across all targets, weighted by the number of ligands per target. Unsurprisingly, models trained on limited and biased public data, perform much better on similar chemistry. Nonetheless, Nesso-1 surpasses Boltz-2 across all levels of training set similarity while achieving an order of magnitude speed-up.

#### 2.3.4 OpenBind affinity benchmark

Before considering internal assays, we also evaluate Nesso-1 zero-shot on the recent OpenBind affinity benchmark [6]. This includes 494 compounds designed to target the EV-A71 2A protease. The benchmark curators highlight how this is an exceptionally challenging target, driven in part by a significant apo-to-holo conformational change where a flexible loop opens to accommodate ligand binding. Beyond AI open-source models (Boltz-2, AQAffinity from SandboxAQ, and AEV-PLIG [60]), they also consider docking-baselines [4] and simple physicochemical ranking scores derived from molecular weight and cLogP, respectively. Following their evaluation, we report RMSE and Spearman *ρ*. We also refer to Figure 1 for the Pearson correlation achieved by both Nesso-1 and Boltz-2 on this benchmark. We emphasize that no additional information about the chemistry or binding site is provided to Nesso-1, so as to allow for a fair comparison with the reported baselines.

##### Analysis of results

Figure 8 shows that Nesso-1 achieves the lowest RMSE and highest Spearman rank correlation. In particular, it convincingly outperforms open-source AI-affinity models, including cofolding-based methods such as Boltz-2, on both metrics. The results are encouraging as this recent dataset offers a more realistic example of an out-of-the-box usage of Nesso-1 on a drug-discovery campaign. To support this point, we refer to Figure 7, which illustrates how the chemical similarity between OpenBind and training set is far lower than that of other benchmarks such as OpenFE and CASP16. However, we note that the difference between Nesso-1 Spearman rank *ρ* and that of molecular weight, is not particularly significant. This underscores that zero-shot evaluation on real-world drug discovery campaigns represents a formidable generalization challenge for models trained exclusively on (biased) public data.

**Figure 8:**
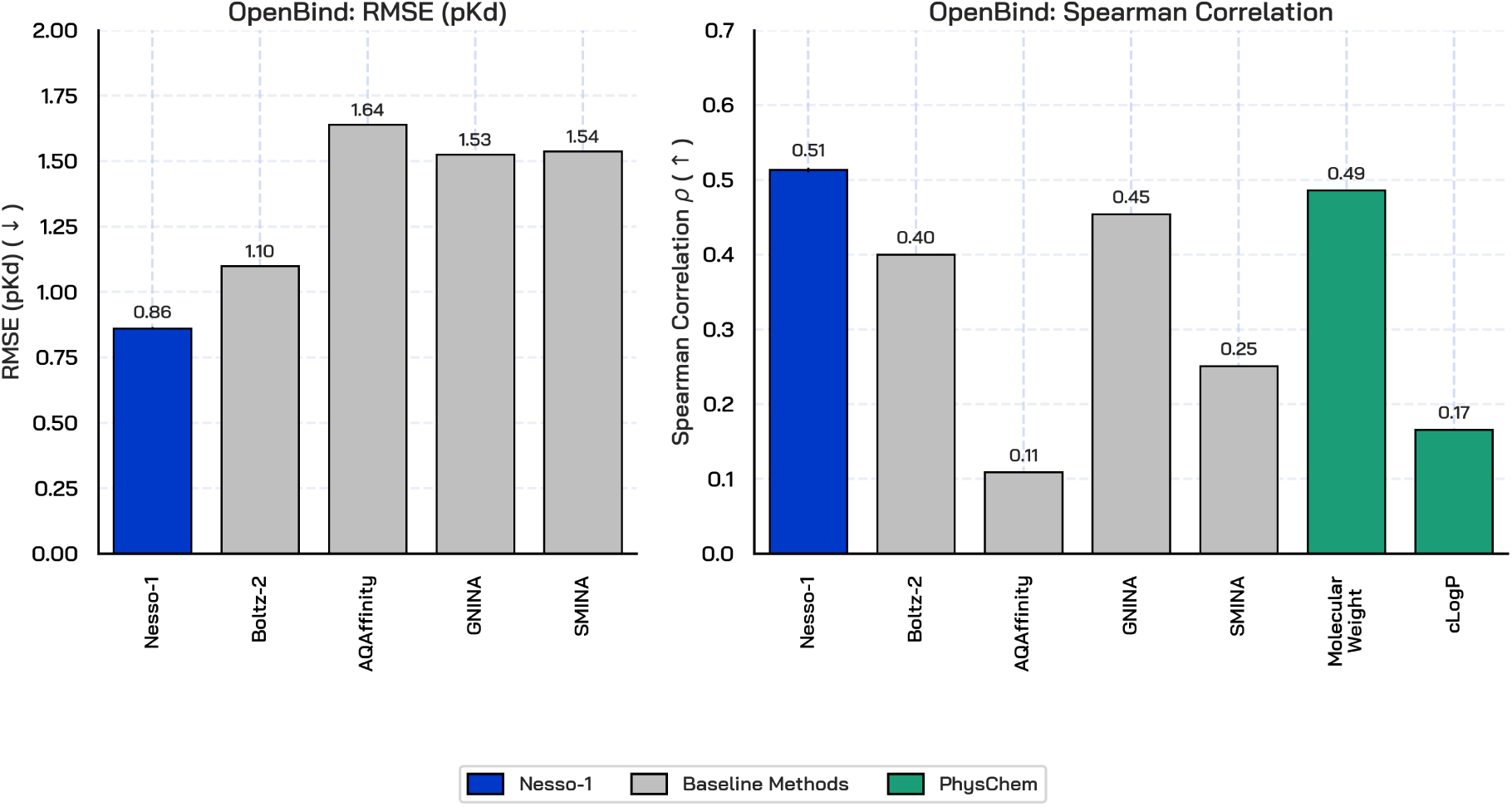
RMSE and Spearman rank of Nesso-1 against AI affinity models, docking scores and physicochemical baselines, evaluated over the recent OpenBind affinity benchmark. Nesso-1 achieves the lowest RMSE and highest Spearman, surpassing other AI baselines by a substantial margin. However, a simple baseline such as molecular weight is the second-best predictor, further highlighting the challenges of zero-shot generalization of models trained on limited (public) bio-chemical space.

#### 2.3.5 Internal biochemical assays

To provide a more complete overview of how Nesso-1 would perform on a drug-discovery campaign, we evaluate it on a selection of 25 internal biochemical assays. These encompass a variety of targets belonging to different protein classes, with an average of several hundred compounds per assay. We primarily benchmark Nesso-1 against the leading open-source affinity model, Boltz-2. In line with the OpenBind benchmark, we also include a molecular-weight ranking, which mainly provides an indirect assessment of assay composition or biases, as across cycles stronger binders may, ultimately, have larger weight. Our evaluation revolves around the following points:

- Show that Nesso-1 outperforms the leading open-source cofolding affinity model (Boltz-2) on real-world medicinal chemistry.
- Demonstrate that Nesso-1*can* remain accurate on assays where molecular weight is not predictive of potency.
- Assess the extent to which Nesso-1 is able to generalize to out-of-distribution chemistry.
- Investigate whether Nesso-1 is *selective* and therefore can be used to detect off-target liabilities.

##### Nesso-1 delivers the highest accuracy

In Figure 9 we compute cross-assay average Pearson Correlation *r*, weighted by the number of compounds in an assay. We compare Nesso-1, Boltz-2—both evaluated *zero-shot*—and molecular weight across the chosen 25 assays. Nesso-1 achieves the highest weighted Pearson correlation. In general, the average performance of Boltz-2 is somewhat comparable to that attained on the OpenBind benchmark, and lower than that reported on other internal assays [34] and public benchmarks [24], suggesting that our selection of assays is, likely, harder for public models such as Boltz-2. We also report the per-assay performance of Nesso-1 and the main baselines in Figure 10. We note that the inherent difficulty of these assays presents a challenge to all baselines evaluated. Consequently, while Nesso-1 maintains a predictive edge over Boltz-2, its absolute performance is lower than that achieved on public benchmarks. Importantly, though, the *>* 10*×* speed-up over Boltz-2 renders these relative accuracy improvements highly impactful in practical applications.

**Figure 9:**
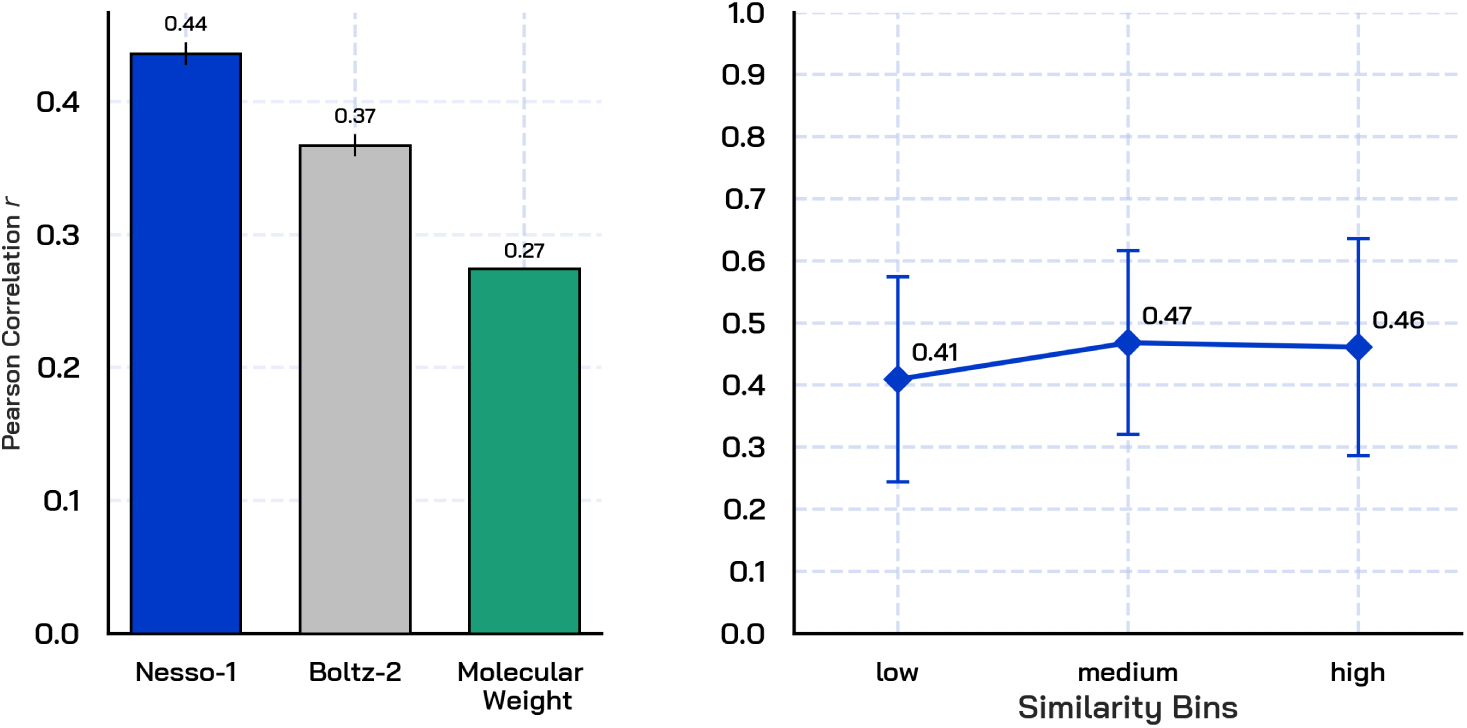
**Left:** Weighted average Pearson Correlation *r* across 25 internal biochemical assays. We compare Nesso-1, Boltz-2 and a molecular-weight ranking baseline. While the absolute performance of the two cofolding models underscores the difficulty of zero-shot generalization to real-world medicinal chemistry, Nesso-1 manages to yield the highest accuracy. **Right:** Weighted average Pearson Correlation attained by Nesso-1 as a function of the internal-assay chemical similarity to the training data. Assay similarity is calculated using Morgan fingerprints and is defined as the mean maximum Tanimoto similarity to the training set across all ligands in an assay. We bin assays from least-similar (low) to most-similar (high). While Nesso-1 is slightly more accurate on the latter, its performance is roughly stable across different levels of chemical similarity.

**Figure 10:**
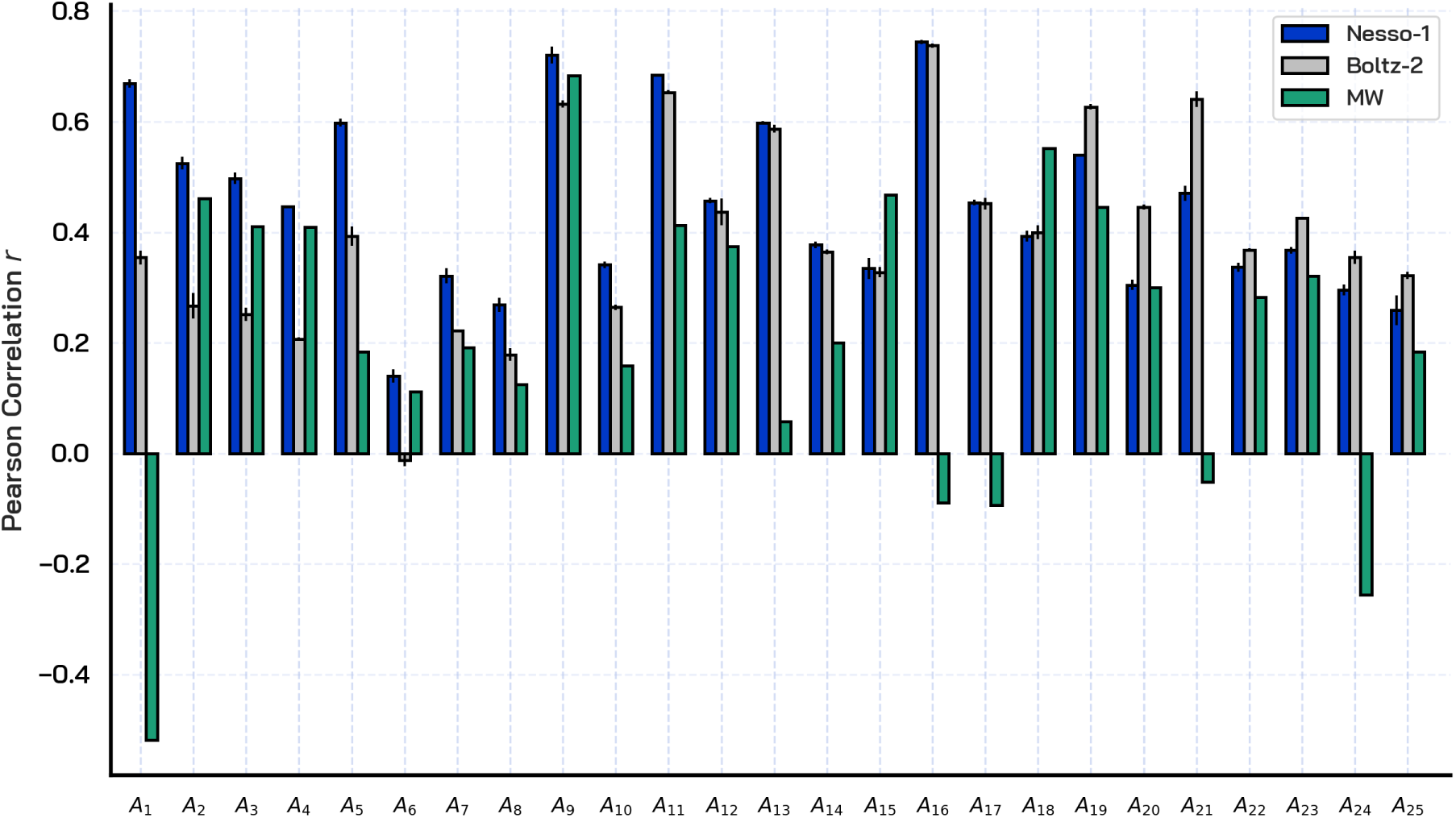
Pearson correlation of Nesso-1, Boltz-2, and a molecular weight baseline evaluated zero-shot on 25 internal biochemical assays. While no model performs well across *all* assays, on average, Nesso-1 achieves the highest Pearson. Importantly, Nesso-1 can remain accurate on most of the assays where molecular weight is not predictive of potency.

##### Nesso-1 is not bound by the predictive power of molecular weight

In general, molecular weight can be a strong baseline (as demonstrated by the OpenBind release) and confirmed by *a few* of our assays. However, it remains an unreliable predictor, as it can fail catastrophically (see for example assays *A*_1_*, A*_16_*, A*_17_*, A*_24_). As such, an ideal cofolding affinity model should be able to deliver good correlation with experiments when molecular weight is poorly predictive of affinity (potency). While there are particularly challenging assays such as *A*_6_ where any baseline struggles, we note that on the aforementioned assays (*A*_1_*, A*_16_*, A*_17_*, A*_24_) where molecular weight is *not* predictive of affinity, both Nesso-1 and Boltz-2 remain accurate. Importantly, on assays *A*_2_*, A*_3_*, A*_4_, Boltz-2 significantly underperforms molecular weight, while Nesso-1 manages to deliver the best correlation.

##### Nesso-1 can achieve good correlation on out-of-distribution assays

Next, in Figure 9 we review how the performance of Nesso-1 correlates with the chemical similarity of each assay to the training set. First, we recall that public benchmarks such as CASP16 and OpenFE are significantly more in-distribution (see Figure 7). Second, we note that while there is an expected trend where lower similarity generally incurs lower accuracy, this is not significant. Nesso-1 remains relatively stable across different levels of chemical similarity of the internal assays. In fact, based on internal evaluations, Nesso-1 can even attain strong correlation on some of the most out-of-distribution assays. This is a promising signal that Nesso-1 is not purely memorizing information from the public data and, thanks to its efficiency, would constitute a meaningful framework to build on. We finally observe that a more comprehensive analysis of the generalization capabilities of Nesso-1 should also encompass the sequence and structure similarity of the internal targets to the training data. However, due to the proprietary nature of the internal datasets, this specific analysis cannot be disclosed in this report.

##### Nesso-1 attains meaningful selectivity on selected assays

A few assays used in our evaluation represent off-targets to a primary target and are hence screened against (subsets of) the same compounds. We investigate whether a cofolding-based approach such as Nesso-1 can correctly identify compounds that are potent for the primary target but not for the off-target—which represent the most plausible drug-candidates. We analyze two different primary targets and their related off-targets in Figure 11 and Figure 12. In particular, we report the pIC50 difference between on- and off-targets for each compound as predicted by Nesso-1 against the experimental potency differences. While there is room for improvement, in the first case, Nesso-1 achieves meaningful selectivity in terms of both Pearson Correlation *r* and MAE, highlighting the benefits of a cofolding-based approach to affinity prediction. In the second case, the overall accuracy is lower, yielding only modest correlation and a higher MAE. This is mainly a consequence of the targets involved, as in the latter case both the on- and off-target are far more out-of-distribution relative to the training data.

**Figure 11:**
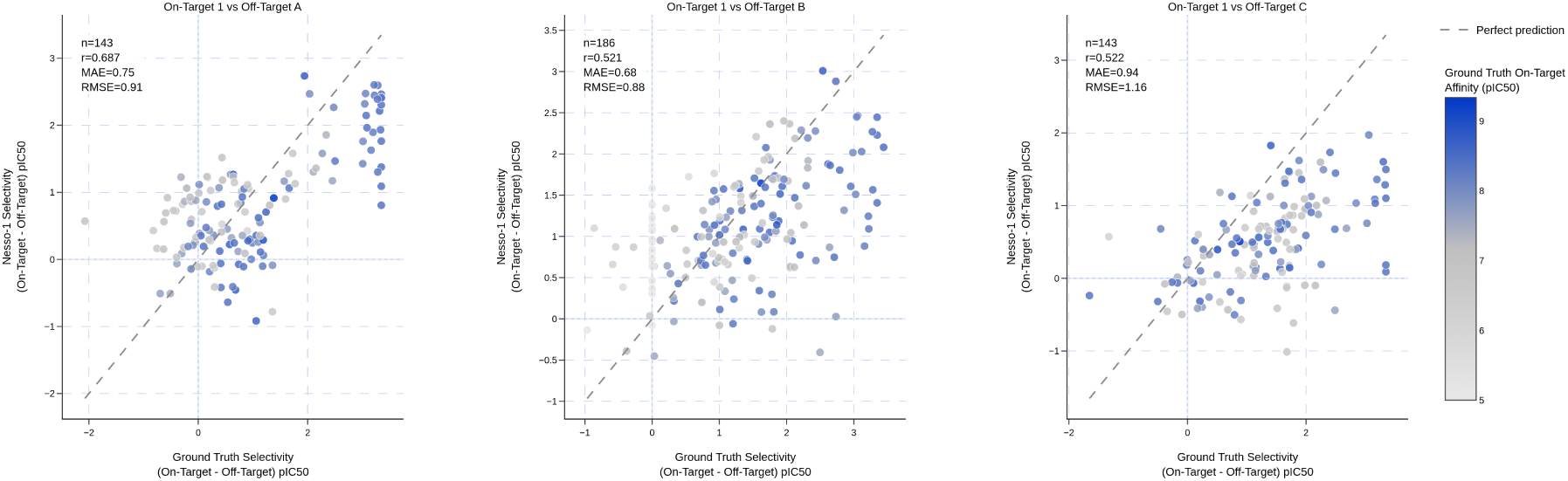
We evaluate Nesso-1 on the same set of compounds screened against a primary target and three related off-targets. Nesso-1 demonstrates meaningful selectivity, achieving decent Pearson Correlation (particularly for Off-Target A) and overall low MAE. The results provide further evidence to the benefit of cofolding-based affinity models.

**Figure 12:**
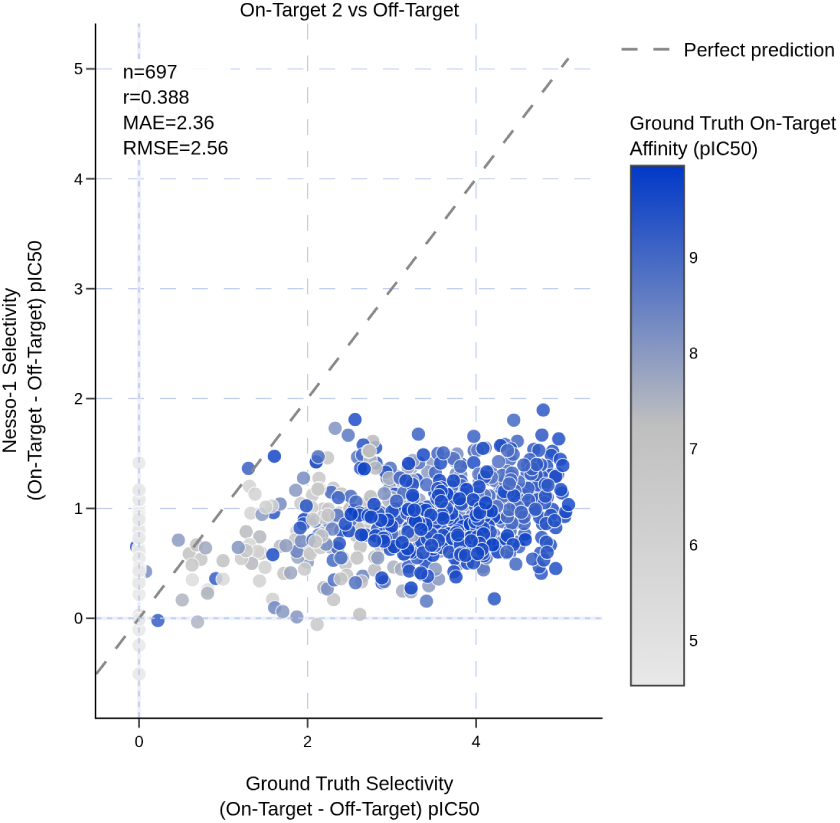
We evaluate Nesso-1 on the same set of compounds screened against a more difficult primary target and related off-target. While Nesso-1 achieves modest correlation, its MAE is high. This is a consequence of the targets involved, as they are far more out-of-distribution for Nesso-1. The results confirm that zero-shot evaluation of public models on under-represented targets and chemistry is an extremely difficult task.

##### Challenges of zero-shot evaluation

The performance across assays reported in Figure 10 further highlights that zero-shot evaluation of models trained on public data over internal, out-of-distribution, medicinal chemistry is an extremely challenging task. Indeed, the accuracy can vary significantly as a function of the target and the chemistry analyzed. We note that internal knowledge of targets and chemistry allows us to draw more meaningful conclusions regarding why certain assays are particularly challenging and where to expect a model like Nesso-1 to perform well. In fact, excluding a priori targets known to be highly out-of-distribution would have inflated our metrics. Nonetheless, we highlight that Nesso-1 manages to outperform the leading open-source cofolding affinity model, Boltz-2. Crucially, the acceleration offered by Nesso-1 enables a broader exploration of the chemical space and significantly facilitates fine-tuning, establishing it as a valuable foundation for the community to build upon.

## 3 Conclusions

In this report we have introduced Nesso-1, a coarse-grained cofolding model for predicting binding affinity. Nesso-1 is more than 10*×* faster than the leading open-source baseline, Boltz-2, with a speed-up of over 20*×* for larger targets, making pre-training and fine-tuning on large libraries extremely more feasible. Crucially, our evaluations across public and internal biochemical assays demonstrate that Nesso-1 advances the state-of-the-art for open-source cofolding affinity models in terms of both accuracy and speed. We illustrate that Nesso-1 can remain competitive on assays with extremely low chemical similarity to the training data or where physicochemical descriptors like molecular weight cannot reliably predict potency. This validates that Nesso-1 has learned meaningful representations that can transfer out-of-distribution. In general, our empirical results suggest that reducing the resolution of affinity models not only diminishes their computational cost, thereby broadening their range of applicability, but may also improve their accuracy over atomistic baselines. We hypothesize that existing approaches such as Boltz-2 struggle to learn meaningful interactions with full heavy-atom resolution. As such, high-frequency information provided by the 3D poses may introduce noise in the predictions and hence hurt performance. We release Nesso-1’s weights to provide to the community a faster affinity model that matches or surpasses that accuracy of leading open-source models.

### Limitations and future updates

While the coarse-graining framework at the core of Nesso-1 allows for fast binding affinity predictions, the lack of atomistic resolution may limit its applicability in structural tasks. Indeed, even though coarse-grained poses may be generated from the trunk output as in [34], Nesso-1 is, primarily, an affinity model. As such, we have focused our evaluation and testing on affinity prediction. Full-atom resolution may still be necessary for tasks where fine-grained physical interactions are paramount. As an orthogonal effort, we note a concurrent open-source release that instead aimed at replicating the structural prediction results of IsoDDE by staying full-atom and expanding the trunk capacity further [61]. However, the precision gained in structure prediction comes at the expense of higher inference costs, further reducing the applicability of these models in early-stages of drug-discovery, where operating across vast regions of chemical space is required. Conversely, Nesso-1 prioritizes speed for binding-affinity prediction by foregoing all-atom resolution. In our report, we show that this design choice not only enables the screening of up to 20*×* the number of compounds using the same compute budget, but also delivers higher accuracy than existing baselines.

Zero-shot evaluations on out-of-distribution benchmarks such as OpenBind and our selection of internal assays, indicate significant potential for further improvement. While Nesso-1’s accuracy could be substantially enhanced by leveraging proprietary ligand embeddings and fine-tuning on internal data, such optimizations fall outside the scope of this release. Our primary objective is to open-source a model trained strictly on public data, deliberately excluding the predictive advantages of exposure to proprietary chemistry and targets. Crucially, Nesso-1 is faster than existing baselines and an overall smaller model, making it a solid starting point to iterate on to further close the performance gap between in-distribution and out-of-distribution medicinal chemistry. Indeed, deployment of Nesso-1 in high-throughput virtual screening, fine-tuning over large data libraries, and extraction of representations for various downstream tasks [62] are all more feasible and far less computationally expensive than relying on all-atom baselines.

More generally, predicting binding affinity through cofolding discards some degrees of freedom, for example involving entropy and water contributions. In fact, during later optimization stages, throughput is no longer the primary objective and a fundamentally different approach to the problem may be required, where all degrees of freedom need to be modeled. We refer to our novel framework AquaGen [63], a generative-model engine that operates at *exactly* the same resolution as Molecular Dynamics as a first attempt at this.

## Acknowledgment

We thank Gail Bartlett, Alan Bilsland, Michael Bronstein, Cristian Gabellini, Daniel Cohen, Adam Kecskes, Ben Mabey, Stephen MacKinnon, Lukáš Pravda, Allison Cohen and Cas Wognum for invaluable feedback and discussion on the manuscript and around this work. We particularly thank Cristian Gabellini for his contributions to earlier versions of the model. We also express our gratitude to Kian Kenyon-Dean, Dragos Cristian Manta, Ihor Neporozhnii, Fereshteh Shakeri, Austin Tripp, Caden Ellis, Aleksandar Djuric, Vitali Marenny for crucial support on testing Nesso-1 code and weights. Finally, we thank Valence Labs and Recursion, for the support and for providing the environment and resources to pursue this work.

## References

[1] Mark A Murcko. The affinity advantage. Journal of Medicinal Chemistry, 69(3):1963–1969, 2026.

[2] Oleg Trott and Arthur J Olson. Autodock vina: improving the speed and accuracy of docking with a new scoring function, efficient optimization, and multithreading. Journal of computational chemistry, 31(2):455–461, 2010.

[3] Richard A Friesner, Jay L Banks, Robert B Murphy, Thomas A Halgren, Jasna J Klicic, Daniel T Mainz, Matthew P Repasky, Eric H Knoll, Mee Shelley, Jason K Perry, et al. Glide: a new approach for rapid, accurate docking and scoring. 1. method and assessment of docking accuracy. Journal of medicinal chemistry, 47(7):1739–1749, 2004.

[4] Andrew T McNutt, Paul Francoeur, Rishal Aggarwal, Tomohide Masuda, Rocco Meli, Matthew Ragoza, Jocelyn Sunseri, and David Ryan Koes. Gnina 1.0: molecular docking with deep learning. Journal of cheminformatics, 13(1):43, 2021.

[5] Edward B Miller, Robert B Murphy, Daniel Sindhikara, Kenneth W Borrelli, Matthew J Grisewood, Fabio Ranalli, Steven L Dixon, Steven Jerome, Nicholas A Boyles, Tyler Day, et al. Reliable and accurate solution to the induced fit docking problem for protein–ligand binding. Journal of Chemical Theory and Computation, 17(4):2630–2639, 2021.

[6] OpenBind Consortium. OpenBind Structure–Affinity Data Release: Enterovirus A71 2A Protease, May 2026.

[7] William L Jorgensen and C Ravimohan. Monte carlo simulation of differences in free energies of hydration. The Journal of chemical physics, 83(6):3050–3054, 1985.

[8] Lingle Wang, Yujie Wu, Yuqing Deng, Byungchan Kim, Levi Pierce, Goran Krilov, Dmitry Lupyan, Shaughnessy Robinson, Markus K Dahlgren, Jeremy Greenwood, et al. Accurate and reliable prediction of relative ligand binding potency in prospective drug discovery by way of a modern free-energy calculation protocol and force field. Journal of the American Chemical Society, 137(7):2695–2703, 2015.

[9] Robert Abel, Lingle Wang, Edward D Harder, BJ Berne, and Richard A Friesner. Advancing drug discovery through enhanced free energy calculations. Accounts of chemical research, 50(7):1625– 1632, 2017.

[10] David F Hahn, Christopher I Bayly, Melissa L Boby, Hannah E Bruce Macdonald, John D Chodera, Vytautas Gapsys, Antonia SJS Mey, David L Mobley, Laura Perez Benito, Christina EM Schindler, et al. Best practices for constructing, preparing, and evaluating protein-ligand binding affinity benchmarks [article v1. 0]. Living journal of computational molecular science, 4(1):1497, 2022.

[11] Gregory A Ross, Chao Lu, Guido Scarabelli, Steven K Albanese, Evelyne Houang, Robert Abel, Edward D Harder, and Lingle Wang. The maximal and current accuracy of rigorous protein-ligand binding free energy calculations. Communications Chemistry, 6(1):222, 2023.

[12] Wentao Li, Chao Lu, Meng Wu, Xuan Wu, Yu Xia, Ziqing Zhang, Xingyuan Xu, Youjun Xu, Yang Chen, Chang Han, et al. Physics-based vs ai-based free energy prediction for protein-ligand potency: Public benchmarks and internal project evidence. ChemRxiv, 2026.

[13] Yiqi Chen and Jian Yang. Acceleration of the gromacs free-energy perturbation calculations on gpus. ACS omega, 10(22):22858–22873, 2025.

[14] Stephan Thaler, Zhiyi Wu, William G Glass, Richard T Bradshaw, Gail Bartlett, Prudencio Tossou, and Geoffrey PF Wood. Boltz-abfe: Free energy perturbation without crystal structures. Journal of Chemical Theory and Computation, 22(4):1823–1833, 2026.

[15] John Jumper, Richard Evans, Alexander Pritzel, Tim Green, Michael Figurnov, Olaf Ronneberger, Kathryn Tunyasuvunakool, Russ Bates, Augustin Žıdek, Anna Potapenko, et al. Highly accurate protein structure prediction with AlphaFold. nature, 596(7873):583–589, 2021.

[16] Josh Abramson, Jonas Adler, Jack Dunger, Richard Evans, Tim Green, Alexander Pritzel, Olaf Ronneberger, Lindsay Willmore, Andrew J Ballard, Joshua Bambrick, et al. Accurate structure prediction of biomolecular interactions with AlphaFold 3. Nature, 630(8016):493–500, 2024.

[17] Minkyung Baek, Frank DiMaio, Ivan Anishchenko, Justas Dauparas, Sergey Ovchinnikov, Gyu Rie Lee, Jue Wang, Qian Cong, Lisa N Kinch, R Dustin Schaeffer, et al. Accurate prediction of protein structures and interactions using a three-track neural network. Science, 373(6557):871–876, 2021.

[18] Rohith Krishna, Jue Wang, Woody Ahern, Pascal Sturmfels, Preetham Venkatesh, Indrek Kalvet, Gyu Rie Lee, Felix S Morey-Burrows, Ivan Anishchenko, Ian R Humphreys, et al. Generalized biomolecular modeling and design with rosettafold all-atom. Science, 384(6693):eadl2528, 2024.

[19] Chai Discovery team, Jacques Boitreaud, Jack Dent, Matthew McPartlon, Joshua Meier, Vinicius Reis, Alex Rogozhonikov, and Kevin Wu. Chai-1: Decoding the molecular interactions of life. BioRxiv, pages 2024–10, 2024.

[20] Jarren Zhuoran Qiao, Feizhi Ding, Thomas Dresselhaus, Mia Rosenfeld, Xiaotian Han, Owen Howell, Aniketh Iyengar, Stephen Opalenski, Anders Christensen, Sai Krishna Sirumalla, et al. Neuralplexer3: accurate biomolecular complex structure prediction with flow models. Advances in Neural Information Processing Systems, 38:1–38, 2026.

[21] Jeremy Wohlwend, Gabriele Corso, Saro Passaro, Noah Getz, Mateo Reveiz, Ken Leidal, Wojtek Swiderski, Liam Atkinson, Tally Portnoi, Itamar Chinn, et al. Boltz-1 democratizing biomolecular interaction modeling. BioRxiv, pages 2024–11, 2025.

[22] Alejandro Dobles, Nina Jovic, Kenneth Leidal, Pranav Murugan, David C Williams, Drausin Wulsin, Nate Gruver, Christina X Ji, Korrawat Pruegsanusak, Gianluca Scarpellini, et al. Pearl: A foundation model for placing every atom in the right location. *arXiv preprint arXiv:2510.24670*, 2025.

[23] ByteDance AML AI4Science Team, Xinshi Chen, Yuxuan Zhang, Chan Lu, Wenzhi Ma, Jiaqi Guan, Chengyue Gong, Jincai Yang, Hanyu Zhang, Ke Zhang, et al. Protenix-advancing structure prediction through a comprehensive AlphaFold3 reproduction. BioRxiv, pages 2025–01, 2025.

[24] Saro Passaro, Gabriele Corso, Jeremy Wohlwend, Mateo Reveiz, Stephan Thaler, Vignesh Ram Somnath, Noah Getz, Tally Portnoi, Julien Roy, Hannes Stark, et al. Boltz-2: Towards accurate and efficient binding affinity prediction. BioRxiv, 2025.

[25] Dominykas Lukauskis, Naail Kashif-Khan, Christopher Tame, and Andrew Potterton. Optimizing molecular glues using free energy perturbation and cofolding methods. ChemRxiv, 2025.

[26] Shunzhou Wan, Xibei Zhang, Xiao Xue, and Peter V Coveney. On the reliability of ai methods in drug discovery: Evaluation of Boltz-2 for structure and binding affinity prediction. *arXiv preprint arXiv:2603.05532,* 2026.

[27] Guillaume Bret, François Sindt, and Didier Rognan. Assessing Boltz-2 performance for the binding classification of docking hits. Journal of Chemical Information and Modeling, 66(3):1511–1521, 2026.

[28] Venkata Sai Sreyas Adury, Pratyush Tiwary, Xinyu Gu, and Mrinal Shekhar. Early-enrichment hit discovery via reversible-work c (t) estimation in metadynamics (ctmd). bioRxiv, pages 2026–02, 2026.

[29] Maxwell Kleinsasser, Brayden J Halverson, Edward Kraft, Sean Francis-Lyon, Sarah E Hugo, Mackenzie R Roman, Ben Miller, Andrew D Blevins, and Ian K Quigley. Hermes: Large del datasets train generalizable protein-ligand binding prediction models. *arXiv preprint arXiv:2602.13503*, 2026.

[30] Kairi Furui and Masahito Ohue. Boltzina: Efficient and accurate virtual screening via docking-guided binding prediction with Boltz-2. *arXiv preprint arXiv:2508.17555*, 2025.

[31] Jiaqi Guan, Wesley Wei Qian, Xingang Peng, Yufeng Su, Jian Peng, and Jianzhu Ma. 3D equivariant diffusion for target-aware molecule generation and affinity prediction. In 11th International Conference on Learning Representations, ICLR 2023, 2023.

[32] Julian Cremer, Tuan Le, Frank Noé, Djork-Arné Clevert, and Kristof T Schütt. Pilot: equivariant diffusion for pocket-conditioned de novo ligand generation with multi-objective guidance via importance sampling. Chemical Science, 15(36):14954–14967, 2024.

[33] Julian Cremer, Tuan Le, Mohammad M Ghahremanpour, Emilia Slugocka, Filipe Menezes, and Djork-Arné Clevert. Flowr. root: a flow matching based foundation model for joint multi-purpose structure-aware 3d ligand generation and affinity prediction. *arXiv preprint arXiv:2510.02578*, 2025.

[34] Matteo Rossi, Ryan Pederson, Miles Wang-Henderson, Ben Kaufman, Edward C Williams, Carl Underkoffler, Owen Lewis Howell, Adrian Layer, Stephan Thaler, Narbe Mardirossian, et al. Terrabind: Fast and accurate binding affinity prediction through coarse structural representations. *arXiv preprint arXiv:2602.07735*, 2026.

[35] Jeffrey Ouyang-Zhang, Pranav Murugan, Daniel J Diaz, Gianluca Scarpellini, Richard Strong Bowen, Nate Gruver, Adam Klivans, Philipp Krähenbühl, Aleksandra Faust, and Maruan Al-Shedivat. Triangle multiplication is all you need for biomolecular structure representations. *arXiv preprint arXiv:2510.18870*, 2025.

[36] Salvatore Candido, Thomas Hayes, Alexander Derry, Roshan Rao, Zeming Lin, Robert Verkuil, Bryan Z Wu, Jin Sub Lee, Elise S Bruguera, Jehan A Keval, et al. Language modeling materializes a world model of protein biology. bioRxiv, pages 2026–06, 2026.

[37] Zeming Lin, Halil Akin, Roshan Rao, Brian Hie, Zhongkai Zhu, Wenting Lu, Nikita Smetanin, Robert Verkuil, Ori Kabeli, Yaniv Shmueli, et al. Evolutionary-scale prediction of atomic-level protein structure with a language model. Science, 379(6637):1123–1130, 2023.

[38] Protein data bank: the single global archive for 3d macromolecular structure data. Nucleic acids research, 47(D1):D520–D528, 2019.

[39] Mihaly Varadi, Damian Bertoni, Paulyna Magana, Urmila Paramval, Ivanna Pidruchna, Malarvizhi Radhakrishnan, Maxim Tsenkov, Sreenath Nair, Milot Mirdita, Jingi Yeo, et al. AlphaFold Protein Structure Database in 2024: providing structure coverage for over 214 million protein sequences. Nucleic acids research, 52(D1):D368–D375, 2024.

[40] Pablo Lemos, Zane Beckwith, Sasaank Bandi, Maarten Van Damme, Jordan Crivelli-Decker, Benjamin J. Shields, Thomas Merth, Punit K Jha, Nicola De Mitri, Tiffany Callahan, AJ Nish, Paul Abruzzo, Romelia Salomon-Ferrer, and Martin Ganahl. SAIR: Enabling deep learning for protein-ligand interactions with a synthetic structural dataset. In The Fourteenth International Conference on Learning Representations, 2026.

[41] Tiqing Liu, Yuhmei Lin, Xin Wen, Robert N Jorissen, and Michael K Gilson. Bindingdb: a web-accessible database of experimentally determined protein–ligand binding affinities. Nucleic acids research, 35(suppl 1):D198–D201, 2007.

[42] Barbara Zdrazil, Eloy Felix, Fiona Hunter, Emma J Manners, James Blackshaw, Sybilla Corbett, Marleen De Veij, Harris Ioannidis, David Mendez Lopez, Juan F Mosquera, et al. The chembl database in 2023: a drug discovery platform spanning multiple bioactivity data types and time periods. Nucleic acids research, 52(D1):D1180–D1192, 2024.

[43] Sunghwan Kim, Jie Chen, Tiejun Cheng, Asta Gindulyte, Jia He, Siqian He, Qingliang Li, Benjamin A Shoemaker, Paul A Thiessen, Bo Yu, et al. Pubchem 2023 update. Nucleic acids research, 51(D1):D1373–D1380, 2023.

[44] Fabian Offensperger, Gary Tin, Miquel Duran-Frigola, Elisa Hahn, Sarah Dobner, Christopher W am Ende, Joseph W Strohbach, Andrea Rukavina, Vincenth Brennsteiner, Kevin Ogilvie, et al. Large-scale chemoproteomics expedites ligand discovery and predicts ligand behavior in cells. Science, 384(6694):eadk5864, 2024.

[45] Kevin G Hicks, Ahmad A Cluntun, Heidi L Schubert, Sean R Hackett, Jordan A Berg, Paul G Leonard, Mariana A Ajalla Aleixo, Youjia Zhou, Alex J Bott, Sonia R Salvatore, et al. Protein-metabolite interactomics of carbohydrate metabolism reveal regulation of lactate dehydrogenase. Science, 379(6636):996–1003, 2023.

[46] Peter J Huber. Robust estimation of a location parameter. In Breakthroughs in statistics: Methodology and distribution, pages 492–518. Springer, 1992.

[47] Sheng Xu, Qiantai Feng, Lifeng Qiao, Hao Wu, Tao Shen, Yu Cheng, Shuangjia Zheng, and Siqi Sun. Foldbench: An all-atom benchmark for biomolecular structure prediction. bioRxiv, pages 2025–05, 2025.

[48] Martin Buttenschoen, Garrett M Morris, and Charlotte M Deane. Posebusters: AI-based docking methods fail to generate physically valid poses or generalise to novel sequences. Chemical Science, 15(9):3130–3139, 2024.

[49] Peter Škrinjar, Jérôme Eberhardt, Janani Durairaj, and Torsten Schwede. Have protein-ligand co-folding methods moved beyond memorisation? BioRxiv, pages 2025–02, 2025.

[50] NVIDIA. cuequivariance. https://github.com/NVIDIA/cuEquivariance, 2024.

[51] David Buterez, Jon Paul Janet, Steven J Kiddle, and Pietro Liò. Mf-pcba: Multifidelity high-throughput screening benchmarks for drug discovery and machine learning. Journal of Chemical Information and Modeling, 63(9):2667–2678, 2023.

[52] Min Li, Zhangli Lu, Yifan Wu, and YaoHang Li. BACPI: a bi-directional attention neural network for compound–protein interaction and binding affinity prediction. Bioinformatics, 38(7):1995–2002, 2022.

[53] Mark McGann. Fred pose prediction and virtual screening accuracy. Journal of chemical information and modeling, 51(3):578–596, 2011.

[54] Richard J Gowers, Irfan Alibay, David WH Swenson, Michael M Henry, Benjamin Ries, Hannah M Baumann, James RB Eastwood, Joshua A Mitchell, David Dotson, Joshua T Horton, et al. The open free energy library. Zenodo, 2023.

[55] Zhiyi Wu, Gerhard Konig, Stefan Boresch, and Benjamin P Cossins. Optimizing absolute binding free energy calculations for production usage. Journal of Chemical Theory and Computation, 21(17):8330–8340, 2025.

[56] Michael K Gilson, Jerome Eberhardt, Peter Škrinjar, Janani Durairaj, Xavier Robin, and Andriy Kryshtafovych. Assessment of pharmaceutical protein–ligand pose and affinity predictions in casp16. Proteins: Structure, Function, and Bioinformatics, 94(1):249–266, 2026.

[57] Isomorphic Labs Team. Accurate Predictions of Novel Biomolecular Interactions with IsoDDE, February 2026.

[58] Francesca Deflorian, Laura Perez-Benito, Eelke B Lenselink, Miles Congreve, Herman WT van Vlijmen, Jonathan S Mason, Chris de Graaf, and Gary Tresadern. Accurate prediction of GPCR ligand binding affinity with free energy perturbation. Journal of Chemical Information and Modeling, 60(11):5563–5579, 2020.

[59] Björn Mattsson and W. Patrick Walters. Identifying and addressing systematic data leakage in protein-ligand affinity benchmarks. BioRxiv, 2026.

[60] Ísak Valsson, Matthew T Warren, Charlotte M Deane, Aniket Magarkar, Garrett M Morris, and Philip C Biggin. Narrowing the gap between machine learning scoring functions and free energy perturbation using augmented data. Communications Chemistry, 8(1):41, 2025.

[61] Aureka AI OpenDDE project. Folding, reasoning, and scaling with open-source drug discovery engine, 2026.

[62] Hyosoon Jang, Hyunjin Seo, Yunhui Jang, Seonghyun Park, and Sungsoo Ahn. Boltz is a strong baseline for atom-level representation learning.

[63] Emmanuel Bengio, Sanjeev Raja, Yui Tik Pang, Kerstin Klaeser, Cristian Gabellini, Nikhil Shenoy, Francesco Di Giovanni, and Prudencio Tossou. Aquagen: Scaling generative models to molecular dynamics precision on thousands of atoms. *arXiv preprint arXiv:2607.03513,* 2026.

